## supplementary_information for "The birth of a bacterial tRNA gene by large-scale, tandem duplication events"

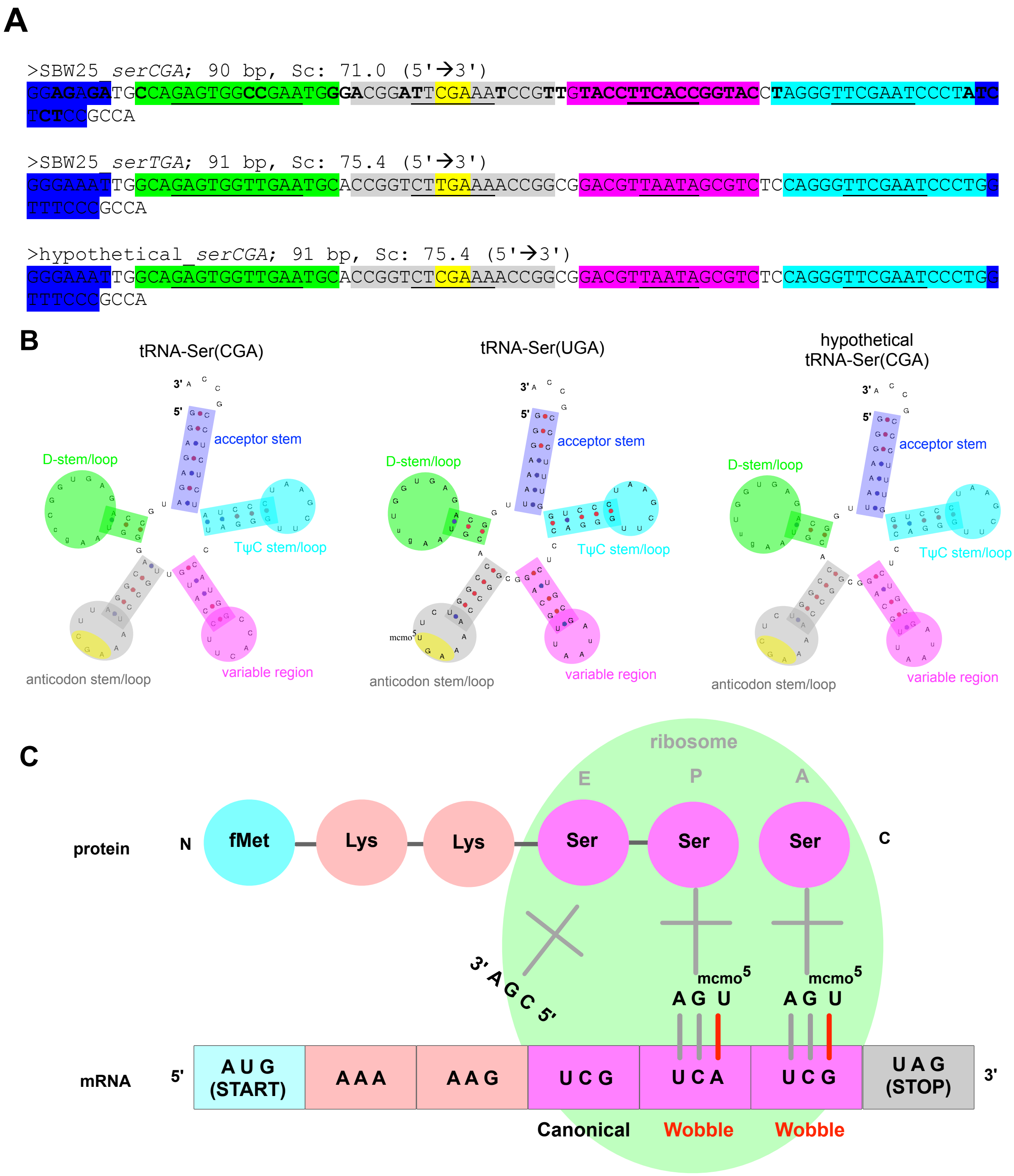


**Figure 1 – supplemental figure 1. Predicted structure and function of tRNA types tRNA-Ser(CGA) and tRNA-Ser(UGA) in *P. fluorescens* SBW25.** (**A**) Gene sequences for *P. fluorescens* SBW25 *serCGA* (top), *P. fluorescens* SBW25 *serTGA* (middle), and hypothetical *serCGA* (*serTGA* backbone with a single point mutation to a CGA anticodon; see Discussion). Highlighting indicates the various parts of each tRNA, with the base=pairing (stem) components in each region underlined (blue=acceptor stem, green=D-stem/loop, grey=anticodon stem/loop, yellow=anticodon, pink=variable region, turquoise=TψC stem/loop). Bold letters in *serCGA* indicate 36 nucleotide differences from *serTGA*. (**B**) Predicted mature tRNA structures for the tRNA sequences in panel A, using tRNAscan-SE v2.0 (Chan and Lowe, 2019). The predicted CmoA/B/M mediated post-transcriptional modification $U_{34}$→mcmo$^5U_{34}$, is also shown in tRNA-Ser(UGA). (**C**) Cartoon depicting translation of serine codons: canonical base pairing occurred between tRNA-Ser(CGA) and codon UCG, and wobble base pairing is occurring between post-transcriptionally modified tRNA-Ser(UGA) and codons UCA and UCG.

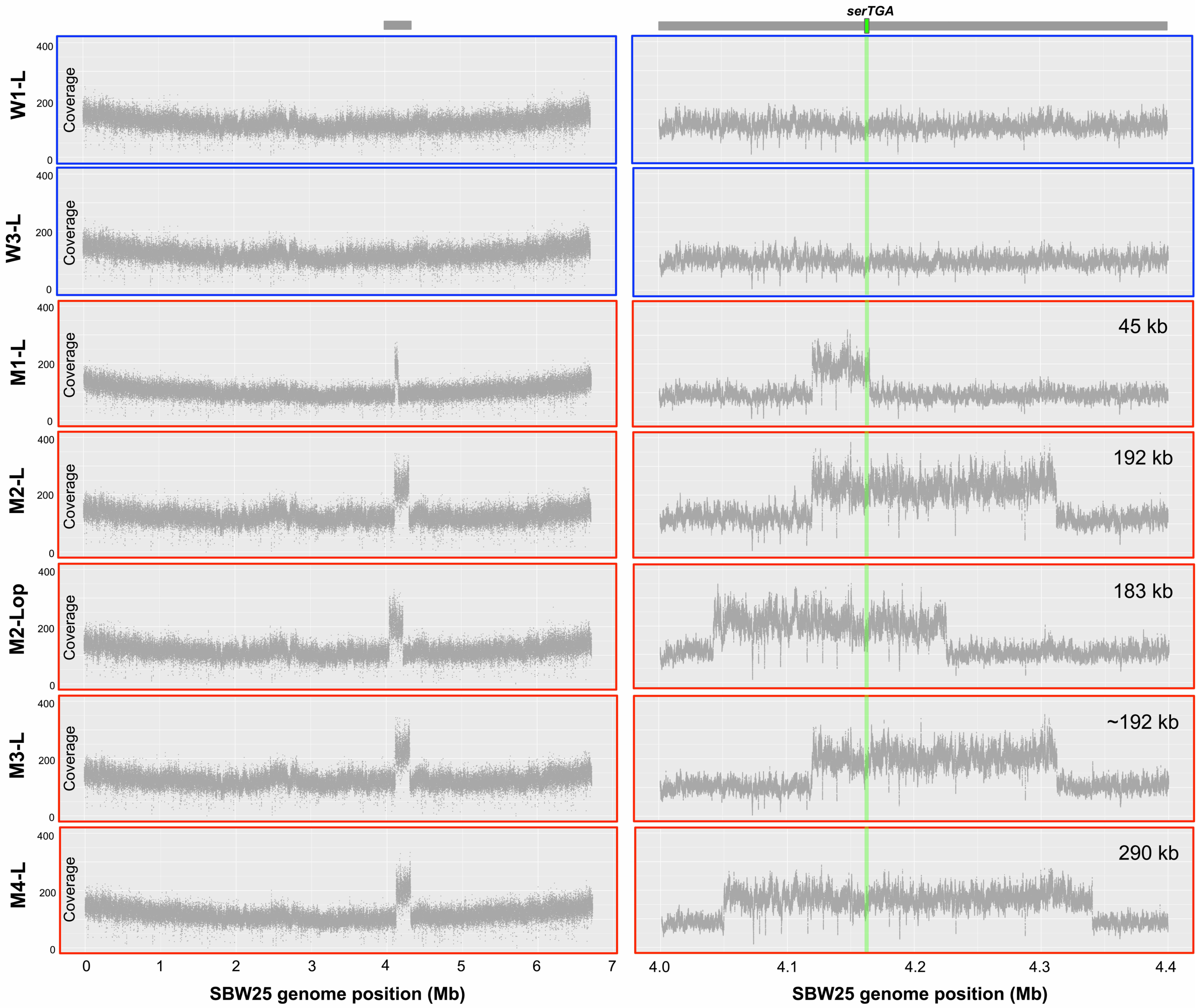


**Figure 4 – supplemental figure 1. Coverage plots from whole genome sequencing data provide evidence of large-scale, tandem duplication events in evolutionary lineages M1-M4.** Whole genome sequence data (Illumina NextSeq, 150 bp, paired-end reads) were obtained for seven strains from Day 13 of the evolution experiment: W1-L and W3-L (derived from independent SBW25 and SBW25-eWT control lineages; outlined in blue), and M1-L M2-L, M2-Lop, M3-L, M4-L (derived from four independent *serCGA* deletion lineages; outlined in red). For each of the seven strains, a minimum of 4.5 million raw sequencing reads were aligned to the SBW25 genome sequence (Silby *et al*., 2009) in Geneious, using the settings described in the main manuscript methods. The number of reads aligned to every 100$^{th}$ base of the SBW25 genome was plotted in R (version 3.6.0; left). This revealed a two-fold increase in coverage between ~4 Mb and ~4.4 Mb, indicating the occurrence of a single, tandem duplication in each of the mutant-derived lineages. Coverage plots for every nucleotide between 4.0 – 4.4 Mb of the SBW25 chromosome (right) demonstrate the occurrence of at least four distinct tandem duplications. These range between 45 kb and 290 kb in size, and each includes a copy of the *serTGA* gene, encoding tRNA-Ser(UGA) (green horizontal bar on zoomed in coverage plots).

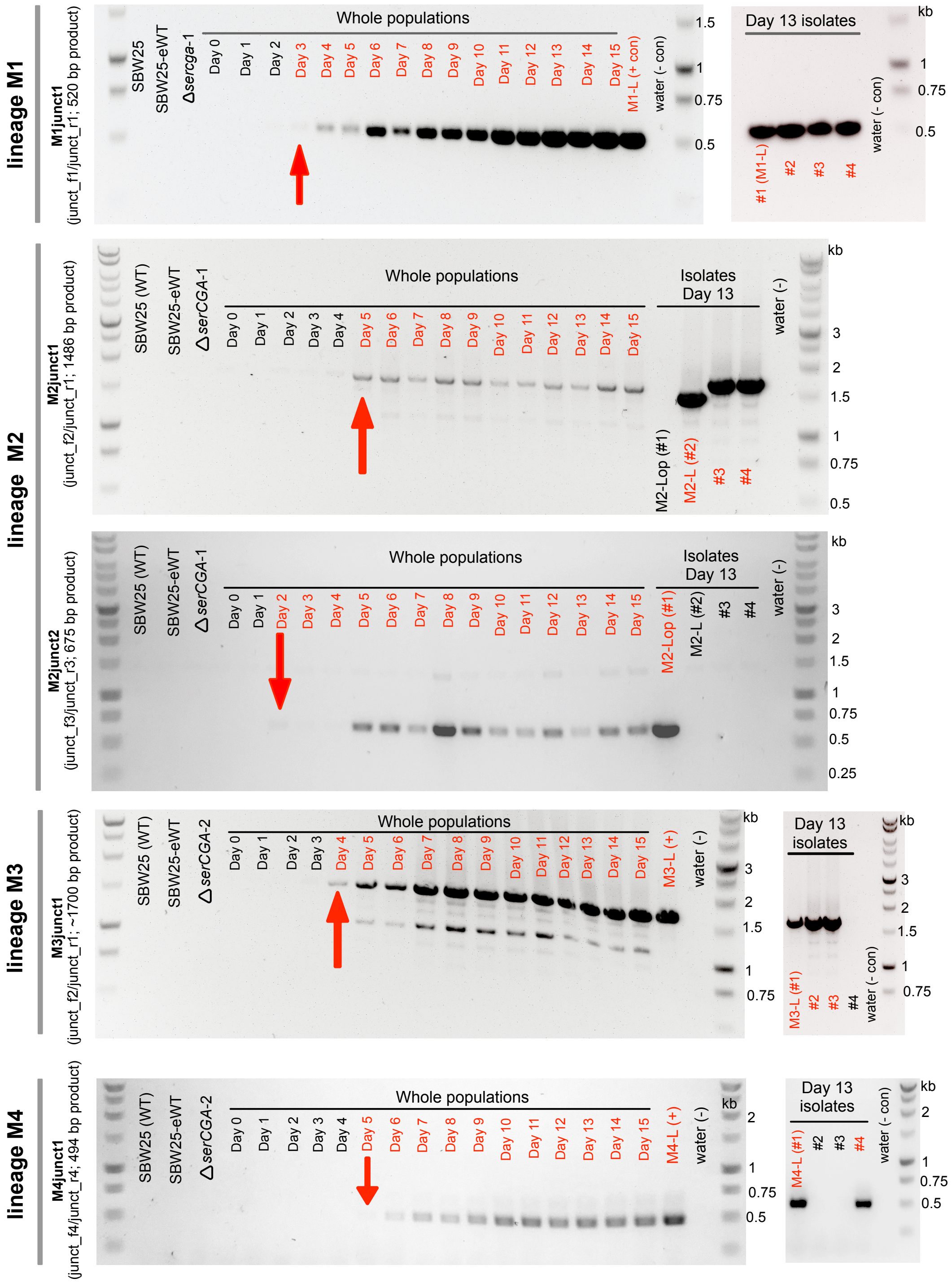


**Figure 4 – supplemental figure 2. Large tandem duplications are first detected between Days 2 and 5 of the evolution experiment.** The emergence of five duplication junctions (top to bottom: M1junct1, M2junct1, M2junct2, M3junct1, M4junct1) was tracked across the relevant lineage (M1, M2, M2, M3, M4, respectively) using duplication junction PCR on daily population samples. Each duplication junction PCR product was first observed between Days 2 and 5 of the evolution experiment (indicated by red arrows). Junction PCRs performed on several individual large colony isolates from each linage indicate that while the duplication fragment identified by whole genome sequencing is present in a significant portion of large colony isolates from each Day 13 population, some genetic heterogeneity exists. Red writing=relevant PCR product detected, black writing=relevant PCR product not detected. The colours in each gel photograph were inverted using Preview (v11.0) to better detect faint OCR products. The left hand gel for lineage M1 is also presented in the main manuscript (Figure 4E), and is included here for completeness.

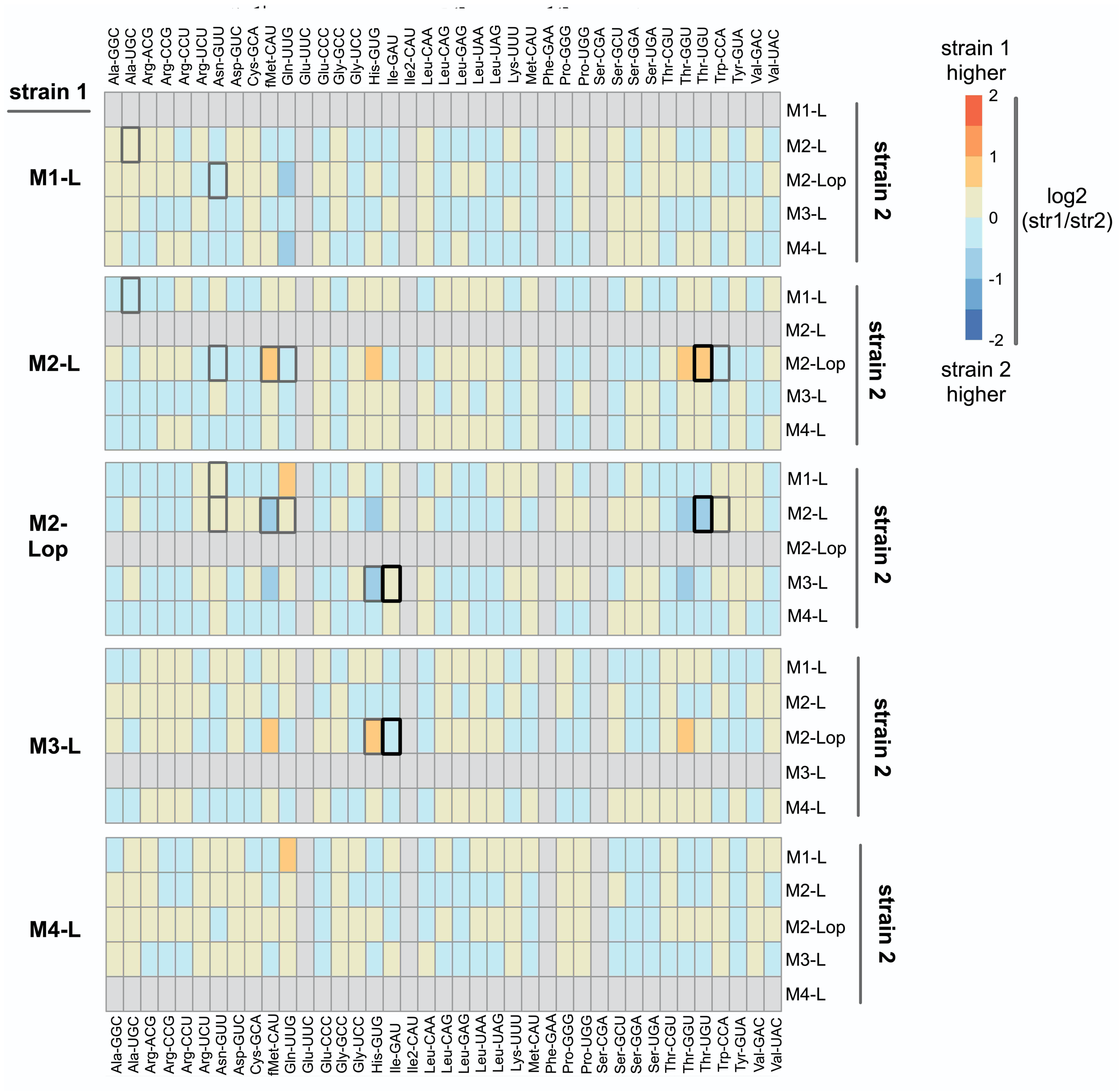


**Figure 5 – supplemental figure 1. Comparison of expression levels of tRNA types in five strains isolated from mutant lineages on Day13.** YAMAT-seq was performed, and data analysed, as described in the manuscript. The read numbers provided in supplemental file 7 were used to make DESeq2 pairwise comparisons between three replicates of each strain (output in source data file 5). Here, the log2-foldchange of each tRNA type is plotted as a heat map for all 20 pairwise comparisons between the five mutant lineage isolates (*e.g.*, M1-L versus each of M2-L, M2-Lop, M3-L and M4-L). Orange/red colouration indicates higher expression in strain 1, blue in strain 2 (colour intensity denotes the size of the difference; darker colour = larger difference). Boxes for three tRNA types (Glu-UUC, Ile2-GAU, Phe-GAA, Ser-CGA) are filled in grey; these were removed from the analysis due to very low YAMAT-seq read numbers for both strains (leading to unreliable DESeq2 output). Boxes for tRNA types with statistically significant differences are highlighted by black or grey borders (black border DESeq2 $p<0.001$, grey $0.001<p<0.01$).

### Reference list for YAMAT-seq: the 42 unique tRNA sequences in *P. fluorescens* SBW25

>1_Ala-GGC-1-2 76 bp Sc: 75.2
GGGGCTATAGCTCAGCTGGGAGAGCGCTTGCATGGCATGCAAGAGGtCAACGGTTCGATCCCGTTTAGCTCCACCA

>2_Ala-TGC-1-5 76 bp Sc: 82.7
GGGGCCATAGCTCAGCTGGGAGAGCGCCTGCCTTGCACGCAGGAGGtCAACGGTTCGATCCCGTTTGGCTCCACCA

>3_Arg-ACG-1-2 77 bp Sc: 84.0
GCACTCGTAGCTCAGCTGGAtAGAGTACTCGGCTACGAACCGAGCGGtCACAGGTTCGAATCCTGTCGAGTGCACCA

>4_Arg-CCG-1-1 77 bp Sc: 83.6
GCATCCGTAGCTCAGCTGGAtAGAGTACTGCCCTCCGAAGGCAGGGGtCGTGGGTTCGAATCCCGCCGGGTGCACCA

>5_Arg-CCT-1-1 77 bp Sc: 72.4
GTCCCAGTAGCTCAATTGGAtAGAGCATCCCCCTCCTAAGGGGAAGGtTGGCCGTTCGAACCGGCCCTGGGACACCA

>6_Arg-TCT-1-1 77 bp Sc: 89.3
GCGCCCGTAGCTCAGCTGGAtAGAGCATCCGCCTTCTAAGCGGATGGtCGCAGGTTCGAGTCCTGCCGGGTGCGCCA

>7_Asn-GTT-1-1 76 bp Sc: 80.6
TCCGTGATAGCTCAGTCGGTAGAGCAAATGACTGTTAATCATTGGGtCCCAGGTTCGAGTCCTGGTCACGGAGCCA

>8_Asn-GTT-2-1 76 bp Sc: 77.6
TCCGCGATAGCTCAGTTGGTAGAGCAAATGACTGTTAATCATTGGGtCCCTGGTTCGAGTCCAGGTCGTGGAGCCA

>9_Asp-GTC-1-4 77 bp Sc: 90.7
GCAGCGGTAGTTCAGTCGGTtAGAATACCGGCCTGTCACGCCGGGGGtCGCGGGTTCGAGTCCCGTCCGCTGCGCCA

>10_Cys-GCA-1-1 74 bp Sc: 65.3
GGCCGAGTAGCAAAATGGTTATGCAGCGGATTGCAAATCCGCCTaCGCCGGTTCGATTCCGACCTCGGCCTCCA

>11_Cys-GCA-2-1 112 bp Sc: 23.4 (pseudo tRNA, no reads align in any sample)
GAGTAAATGTTGGTgAtgcggaatagattatcatataacttattgaaaataatgaggaTATTCGTGGATTGCAAATCCGC
CTaCGCCGGTTCGATTCCGACCTCGGCCTCCA

>12_Gln-TTG-1-1 75 bp Sc: 69.7
AGGGGCGTCGCCAAGCGGTAAGGCAGCAGGTTTTGATCCTGCCATgCGTTGGTTCGAATCCAGCCGCCCCTGCCA

>13_Glu-TTC-1-4 76 bp Sc: 68.9
GTCCCCTTCGTCTAGTGGCctAGGACACCGCCCTTTCACGGCGGTAaCAGGGGTTCGAGTCCCCTAGGGGACGCCA

>14_Gly-CCC-1-1 74 bp Sc: 74.9
GCGGGTATAGTTTAATGGTAGAACAGTAGCTTCCCAAGCTTCCGaCGAGGGTTCGATTCCCTCTACCCGCTCCA

>15_Gly-GCC-1-3 76 bp Sc: 88.4
GCGGGAATAGCTCAGTTGGTAGAGCACGACCTTGCCAAGGTCGGGGtCGCGAGTTCGAGTCTCGTTTCCCGCTCCA

>16_Gly-TCC-1-1 74 bp Sc: 83.2
GCGGGTATAGTTTAGTGGTAGAACCTCAGCCTTCCAAGCTGATGaTGCGGGTTCGATTCCCGCTACCCGCTCCA

>17_His-GTG-1-2 76 bp Sc: 73.3
GTGGGCGTAGCTCAGTTGGTAGAGCACGGGATTGTGACTCCCGTTGtCGAGGGTTCGATCCCCTTCGTCCACCCCA

>18_Ile-GAT-1-5 77 bp Sc: 88.9
GGGTCTGTAGCTCAGTTGGTtAGAGCGCACCCCTGATAAGGGTGAGGtCGGCAGTTCGAATCTGCCCAGACCCACCA

>19_Ile2-CAT-1-1 77 bp Sc: 89.4
GGGCCTATAGCTCAGTTGGTtAGAGCAGGGGACTCATAATCCCTTGGtCGTAGGTTCGAGTCCTACTGGGCCCACCA

>20_Leu-CAA-1-1 85 bp Sc: 70.8

GCCCTGATGGCGGAATTGGTaGACGCGGCGGATTCAAAATCCGTTTTCGAAAgGAGTGGGAGTTCGAGTCTCCCTCGGGG
CACCA

>21_Leu-CAG-1-2 87 bp Sc: 71.6
GCCCAGGTGGTGAAATTGGTaGACACGCCAGCTTCAGGTGCTGGTGATCGCAAGGTCGTGGAAGTTCGAGTCTTCTCCTG
GGCACCA

>22_Leu-GAG-1-1 86 bp Sc: 60.3
GCCGAGGTGGTGGAATTGGTaGACACGCAACCTTGAGGTGGTTGTGCCCATAGGGTgTAGGGGTTCGAGTCCCCTTCTCG
GTACCA

>23_Leu-TAA-1-1 87 bp Sc: 71.0
GCCCGAATGGCGAAACTGGTaGACGCATGGGACTTAAAATCCCCCGCTCGTAAGGGCGTCCCGGTTCGATTCCGGGTTCG
GGCACCA

>24_Leu-TAG-1-1 85 bp Sc: 70.1
GCGGATGTGGTGGAATTGGTaGACACACTGGATTTAGGTTCCAGCGCCGCGAGGCGTAAGAGTTCGAGTCTCTTCATCCG
CACCA

>25_Lys-TTT-1-2 76 bp Sc: 86.5
GGGTCGTTAGCTCAGTTGGTAGAGCAGTTGGCTTTTAACCAATTGGtCGTAGGTTCGAATCCCACACGACCCACCA

>26_Met-CAT-1-1 77 bp Sc: 77.5
GGCTACATAGCTCAGTTGGTtAGAGCATAGCATTCATAATGCTGGGGtCCGGGGTTCAAGTCCCTGTGTAGCCACCA

>27_Phe-GAA-1-1 76 bp Sc: 81.1
GCCCAGATAGCTCAGTCGGTAGAGCAGGGGATTGAAAATCCCCGTGtCGGCGGTTCGATTCCGTCTCTGGGCACCA

>28_Pro-GGG-1-1 77 bp Sc: 67.6
CGGGGCGTAGCGCAGTCcGGTAGCGCACTAGCATGGGGTGCTAGGGGtCGAGTGTTCGAATCACTCCGTCCCGACCA

>29_Pro-TGG-1-2 77 bp Sc: 73.4
CGGGGTATAGCGCAGTCcGGTAGCGCGCCTGCTTTGGGAGCAGGATGtCAGGAGTTCGAATCCCCTTACCCCGACCA

>30_Ser-CGA-1-1 90 bp Sc: 71.0
GGAGAGATGCCAGAGTGGCcgAATGGGACGGATTCGAAATCCGTTGTACCTTCACCGGTACCTAGGGTTCGAATCCCTAT
CTCTCCGCCA

>31_Ser-GCT-1-1 91 bp Sc: 75.0
GGAGAGCTGGCCGAGTGGCcgAAGGCGCTCCCCTGCTAAGGGAGTACACCTCAAaAGGGTGTCGGGGGTTCGAATCCCCC
GTTCTCCGCCA

>32_Ser-GGA-1-1 90 bp Sc: 75.7
GGTGAAGTGTCCGAGTGGCttAAGGAGCACGCCTGGAAAGTGTGTATACAAGAAATTGTATCGAGAGTTCGAATCTCTCC
TTCACCGCCA

>33_Ser-TGA-1-1 91 bp Sc: 75.4
GGGAAATTGGCAGAGTGGTtgAATGCACCGGTCTTGAAAACCGGCGGACGTTAAtAGCGTCTCCAGGGTTCGAATCCCTG
GTTTCCCGCCA

>34_Thr-CGT-1-1 73 bp Sc: 69.7
GCCCGTGTAGCTCAGTCGGTAGAGCAGCGCACTCGTAACGCGAAGGtCGCAGGTTCGATTCCTGTCTCGGGCACCA

>35_Thr-GGT-1-1 76 bp Sc: 80.7
GCTCTTGTAGCTCAGTTGGTAGAGCACACCCTTGGTAAGGGTGAGGtCAGCGGTTCAAATCCGCTCAAGAGCTCCA

>36_Thr-TGT-1-1 76 bp Sc: 87.9
GCCGGTATAGCTCAGTTGGTAGAGCAACTGACTTGTAATCAGTAGGtCCCGGGTTCGACTCCTGGTGCCGGCACCA

>37_Trp-CCA-1-1 76 bp Sc: 86.5
AGGTCAGTAGCTCAATTGGCAGAGCGACGGTCTCCAAAACCGTAGGtTGGGGGTTCGATTCCCTCCTGACCTGCCA

>38_Tyr-GTA-1-1 85 bp Sc: 77.1

GGAGGGGTTCCCGAGCGGCcaAAGGGATCAGACTGTAAATCTGACGTCTACGACTtCGAAGGTTCGAATCCTTCCCCCTC
CACCA

>39_Val-GAC-1-1 77 bp Sc: 78.1
AGGCACGTAGCTCAGTTGGTtAGAGCACCACCTTGACATGGTGGGGGtCGTTGGTTCGAGTCCAATCGCGCCTACCA

>40_Val-TAC-1-3 76 bp Sc: 84.4
GGGTGATTAGCTCAGCTGGGAGAGCATCTGCCTTACAAGCAGAGGGtCGGCGGTTCGATCCCGTCATCACCCACCA

>41_fMet-CAT-1-1 77 bp Sc: 78.4
CGCGGGGTGGAGCAGTCtGGTAGCTCGTCGGGCTCATAACCCGAAGGtCGTCGGTTCAAATCCGGCCCCCGCAACCA

>42_fMet-CAT-2-2 77 bp Sc: 76.7
CGCGGGATGGAGCAGTCtGGTAGCTCGTCGGGCTCATAACCCGAAGGtCGTCGGTTCAAATCCGGCTCCCGCAACCA
